# Impaired sleep-dependent memory consolidation predicted by reduced sleep spindles in Rolandic epilepsy

**DOI:** 10.1101/2024.05.16.594515

**Authors:** Hunki Kwon, Dhinakaran M. Chinappen, Elizabeth A. Kinard, Skyler K. Goodman, Jonathan F. Huang, Erin D. Berja, Katherine G. Walsh, Wen Shi, Dara S. Manoach, Mark A. Kramer, Catherine J. Chu

## Abstract

**Background and Objectives:** Sleep spindles are prominent thalamocortical brain oscillations during sleep that have been mechanistically linked to sleep-dependent memory consolidation in animal models and healthy controls. Sleep spindles are decreased in Rolandic epilepsy and related sleep-activated epileptic encephalopathies. We investigate the relationship between sleep spindle deficits and deficient sleep dependent memory consolidation in children with Rolandic epilepsy.

**Methods:** In this prospective case-control study, children were trained and tested on a validated probe of memory consolidation, the motor sequence task (MST). Sleep spindles were measured from high-density EEG during a 90-minute nap opportunity between MST training and testing using a validated automated detector.

**Results:** Twenty-three children with Rolandic epilepsy (14 with resolved disease), and 19 age- and sex-matched controls were enrolled. Children with active Rolandic epilepsy had decreased memory consolidation compared to control children (p=0.001, mean percentage reduction: 25.7%, 95% CI [10.3, 41.2]%) and compared to children with resolved Rolandic epilepsy (p=0.007, mean percentage reduction: 21.9%, 95% CI [6.2, 37.6]%). Children with active Rolandic epilepsy had decreased sleep spindle rates in the centrotemporal region compared to controls (p=0.008, mean decrease 2.5 spindles/min, 95% CI [0.7, 4.4] spindles/min). Spindle rate positively predicted sleep-dependent memory consolidation (p=0.004, mean MST improvement of 3.9%, 95% CI [1.3, 6.4]%, for each unit increase in spindles per minute).

**Discussion:** Children with Rolandic epilepsy have a sleep spindle deficit during the active period of disease which predicts deficits in sleep dependent memory consolidation. This finding provides a mechanism and noninvasive biomarker to aid diagnosis and therapeutic discovery for cognitive dysfunction in Rolandic epilepsy and related sleep activated epilepsy syndromes.

## Introduction

Rolandic epilepsy (RE, also called self-limited epilepsy with centrotemporal spikes, SeLECTS; previously called benign Rolandic epilepsy; childhood epilepsy with centrotemporal spikes; or benign epilepsy with centrotemporal spikes) is a common focal developmental epilepsy that accounts for 8-23% of childhood epilepsy^1,2^. RE is characterized by a transient period of sleep-potentiated seizures and epileptiform discharges best localized to the inferior Rolandic cortex during childhood. Seizures in RE usually present before 11 years old, with a peak at 5-8 years^1^, and always remit by late adolescence, though epileptiform activity can persist for years after seizure resolution^3^. In addition to seizures, children with RE exhibit cognitive deficits during school-age years^4^ that also resolve by a decade after seizure resolution^1^. The severity of cognitive symptoms varies, but approximately 7% of children with RE have severe cognitive deficits consistent with a severe epileptic encephalopathy^5^.

The mechanism of cognitive dysfunction in RE is an area of active research. We recently discovered that epileptiform spikes interfere with sleep spindles, prominent bursts of 9-15 Hz oscillations mechanistically linked to memory consolidation during stage 2 sleep^6,7^, in RE and severe sleep activated epileptic encephalopathies^8-10^. In contrast to measures of spike rate, sleep spindle rate predicts IQ, processing speed, and sensorimotor coordination in RE^8^, and changes in spindle rate predict cognitive response to high-dose benzodiazepine treatment in severe epileptic encephalopathies^9^. Similar findings have subsequently been reported in other epilepsy populations^11,12^, suggesting that sleep spindle deficits may provide a robust, generalizable biomarker for several cognitive symptoms observed in epilepsy.

Sleep spindles have been most strongly linked to the cognitive process of sleep-dependent memory consolidation^13-21^. Whether spindle deficits relate specifically to deficits in sleep-dependent memory consolidation in epilepsy remains unknown. To evaluate for evidence of deficits in sleep-dependent memory consolidation in children with Rolandic epilepsy and investigate the link between this memory function and sleep spindles, we performed a prospective trial in children with RE and typically developing controls. We hypothesized that memory consolidation would be lower in children with active RE and that sleep spindle rate would predict the degree of memory consolidation over the period of sleep recorded. Identifying this cognitive deficit and demonstrating a relationship with sleep spindles provides both a mechanism and noninvasive biomarker for cognitive dysfunction in RE.

## Methods

### Subjects

We performed a prospective, case-control study in children with RE and age- and sex-matched controls. Children with RE were required to have a history of at least one focal motor or generalized seizure and an EEG with sleep-activated centrotemporal spikes. Children with RE were classified as having active epilepsy (seizure within 12 months) or resolved epilepsy (seizure-free for at least 12 months). We chose this classification because most children with RE who are seizure-free for 1 year have entered sustained remission from epilepsy^1^. Control children were required to have no history of seizures and no known neurological disorders. Children with attention disorders and mild learning difficulties were included, as these profiles are consistent with known RE comorbidities^4^. Eligible children were identified from the community and Massachusetts General Hospital (MGH) pediatric neurology and general pediatrics clinics, the EEG lab, and through posted advertisements. Informed consent was obtained from all participants and this study was approved by the Massachusetts General Hospital Institutional Review Board.

### Experimental overview

The experimental timeline is outlined in **Figure 1**. Briefly, the subjects arrived at the Athinoula A. Martinos Center for Biomedical Imaging at approximately 10:00 AM and completed training on the finger tapping motor sequence typing task (MST) on each hand using separate sequences. This was followed by EEG placement and a nap opportunity of approximately 90 minutes. At approximately 3:00 PM, the nap opportunity ended, and the subjects were tested on the MST sequences on each hand. The subjects then underwent a brain MRI in an adjacent imaging suite after a small meal. Data were processed and analyzed as described below.

**Figure 1.**
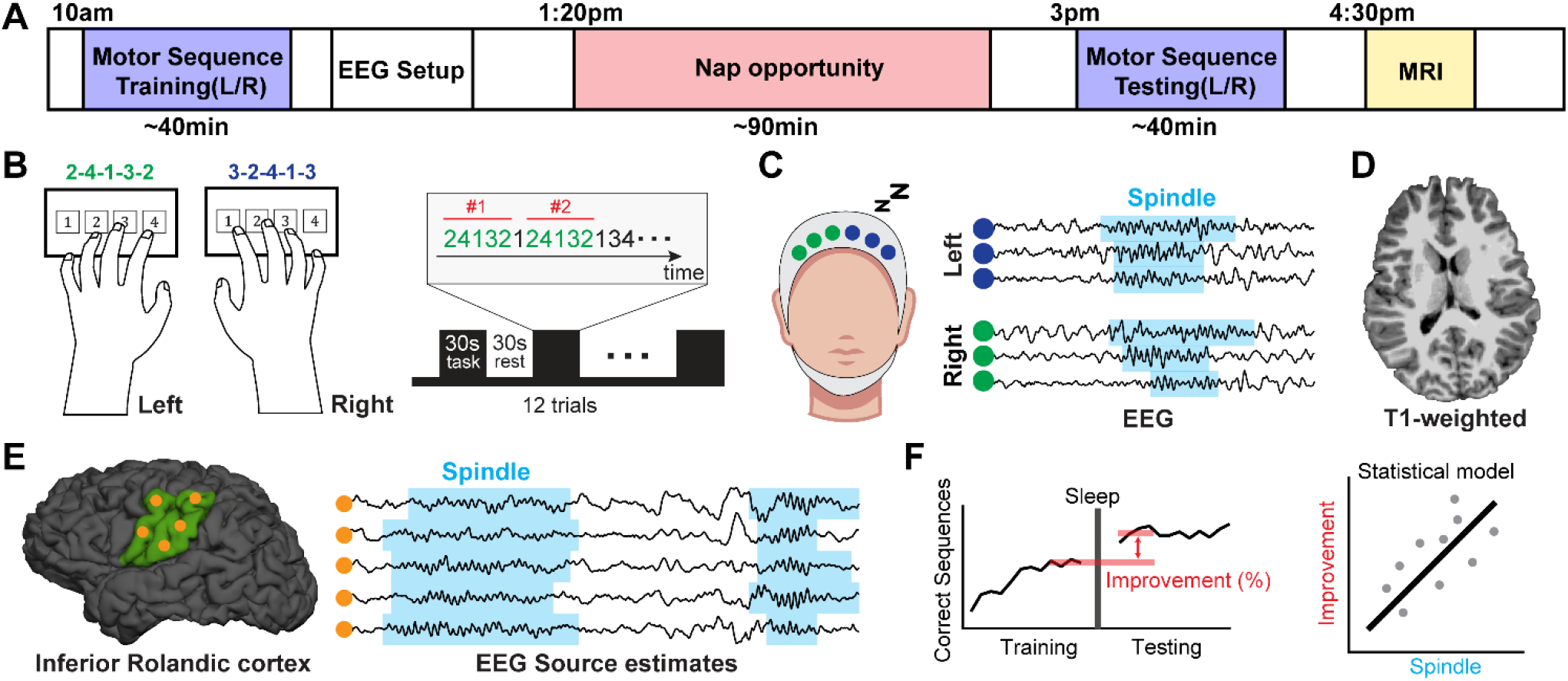
Illustration of the study design. **A)** Overview of the experiment. **B)** Subjects were trained on the motor sequence typing task (MST) with the left and right hands before the nap opportunity and tested after the nap opportunity. **C)** During the nap opportunity, high density EEG were recorded from which sleep spindles were subsequently detected. **D)** After the MST testing, an MRI was obtained. **E)** Sleep spindles were also detected in the inferior Rolandic cortex using electrical source imaging. **F)** Memory consolidation was measured from the MST task and compared to spindle rate during the nap opportunity.

### Finger tapping motor sequence task

For our memory task, we used a well-validated probe of sleep dependent memory consolidation, the finger tapping motor sequence task (MST). This task has been validated to capture sleep-dependent memory consolidation in healthy adults, where performance improves after sleep compared to an equal period of wakefulness^22^.

Subjects were asked to place their fingers on four numerically labeled keys on a standard number pad. They were then instructed to repeatedly type a 5-digit sequence (e.g., 4-1-3-2-4) as quickly and accurately as possible during twelve 30 second trials separated by 30 second rest periods (Figure 1B)^22^. To minimize the involvement of working memory, the sequence was displayed on a monitor during the typing trials. Subjects were trained on separate sequences with their left and right hands before sleep, with a 10-minute break between left- and right-hand training sessions. Following sleep, subjects were tested on the same sequences on the left and right hand respectively with an intervening 10-minute break.

To exclude performance outliers across trials, for each subject and each hand, we fit an exponential model to the learning curve during training, and included a constant offset to the exponential model during the post-sleep testing, as in ^23^. If performance on a trial was less than expected (defined as more than two standard deviations below the model fit), it was excluded as an outlier. The causes of outliers include inattention, task interruption, or misplacement of the fingers on the keys, for example.

To quantify sleep-dependent memory consolidation, for each hand, we calculated the percentage difference between the mean of the last three correct sequences in the training session and the mean of the first three correct sequences in the testing session, following previously published procedures^22^.

### EEG recordings

EEG recordings were acquired with a 70-channel cap (Easycap, Vectorview, Elekta-Neuromag, Helsinki, Finland) at a sampling rate of 2035 Hz or 2000 Hz, and the data were subsequently downsampled to 407 Hz or 400 Hz for analysis. In each case, impedances were maintained below 10 kΩ. Electrode locations were digitized using a 3D digitizer (Fastrak, Polhemus Inc., Colchester, VA). All recordings were manually inspected and sleep-staged by a board-certified neurophysiologist following standard procedures. To target sleep spindles, N2 was selected for analysis, as done previously. Channels with continuous artifact were excluded. The remaining data were re-referenced to an average signal for subsequent analysis.

### MRI recordings

Following the MST testing, all subjects underwent a same-day brain MRI. MRI data were collected on a 3T Magnetom Prisma Siemens MRI scanner with a 64-channel head coil at the Martinos Center for Biomedical Imaging with the following parameters: Multiecho MEMPRAGE (TE = 1.74 milliseconds, TR = 2530 milliseconds, flip angle = 7°, voxel size = 1 × 1 × 1 mm^3^), T2 FLAIR (TE = 3 milliseconds, TR = 5000 milliseconds, flip angle = 7°, voxel size = 0.9×0.9×0.9 mm^3^). ESI analysis was performed with the MNE-C software package (https://mne.tools/stable/index.html/). Cortical brain surfaces were reconstructed using FreeSurfer 7.1.1 (https://surfer.nmr.mgh.harvard.edu/) from MEMPRAGE and T2 FLAIR data.

### Electrical Source Imaging (ESI)

Digitized electrode coordinates were aligned to the MEMPRAGE data using the nasion and auricular points as fiducial markers, and multiple points collected along the facial surface. A three-compartment boundary element model was generated for the forward model using the watershed algorithm in FreeSurfer^24^. EEG source activity was estimated using the dSPM algorithm in the MNE software package^25^. To do so, source estimates of EEG activity were generated for 10,242 vertices per hemisphere. Then, source activity was averaged across 162 vertices per hemisphere by averaging across circles with 1 cm radius using a full-width half-max smoothing kernel in MNE.

Because epileptiform spikes in RE have been well-localized to the inferior Rolandic cortex, and we have previously reported a spindle deficit in this region in a separate cohort^26^, this cortical region was selected as our Region of Interest (ROI) for source analysis. To define this ROI, for each subject, a sphere from the most superior vertex in the Rolandic cortex with a radius equal to half of the distance between the most superior and inferior vertices in Rolandic cortex was labeled and excluded from the precentral and postcentral gyrus labels in Freesurfer, following procedures described previously^26,27^.

### Sleep spindle detector

To measure sleep spindles, we applied an automated spindle detector that we developed specifically to perform well in the setting of sharp events, which are common features in the EEG recordings from patients with epilepsy and young children^8^. To build a spindle detector robust to both healthy and disease states, the detector was trained and validated on manual spindle markings from healthy children, children with continuous spike and wave sleep with encephalopathy, and children with Rolandic epilepsy, resulting in 19,625 manually marked spindles from 115 unique pediatric subjects^28^ not included in this project.

To apply the spindle detector, for each EEG channel or source space time series of interest, we evaluated 0.5 s intervals of data and computed three features: theta band power (4-8 Hz), sigma band power (9-15 Hz), and the Fano factor of the oscillation intervals, which is a measure of cycle regularity^8^. We chose 0.5 s intervals, which are the typical minimum duration accepted for sleep spindles^29^, to maintain a 2 Hz frequency resolution, allowing reliable estimation of the theta band power. Spindle detections separated by less than 1 s were concatenated^8^.

For EEG data, spindle rate was averaged across electrodes in each brain region (Central, Temporal, Frontal, Parietal and Occipital) as in ^8^. For ESI data, the spindle rate across all vertices within the inferior Rolandic cortex were averaged to create a single spindle rate for this ROI per hemisphere, as in ^26^.

### Statistical analyses

To test for differences in spindle rate between groups (control, resolved RE, and active RE), we applied a mixed-effects linear model with spindle rate as the dependent variable, group and age as predictors, and a random subject-specific intercept to account for two observations per subject (left and right hemispheres for each subject). For our primary hypothesis that spindle rate is decreased in the epileptogenic region in Rolandic epilepsy, this model was estimated from spindle rates recorded in the centrotemporal scalp and inferior Rolandic cortex. To confirm the spatial specificity of this result, this model was also applied to other brain regions.

To test for evidence of sleep-dependent memory consolidation using the MST, we applied a mixed-effects linear model with sleep-dependent memory consolidation (see ***Data analysis)*** as the dependent variable; age, and whether the subject slept or remained awake during the rest opportunity as predictors; and a subject-specific intercept to account for two observations per subject (improvement from the left and right hands for each subject). We estimated the model separately for each patient group.

To test for differences in sleep-dependent memory improvement between groups for subjects who slept during the rest opportunity, we estimated a mixed-effects linear model with sleep-dependent memory consolidation as the dependent variable, group and age as predictors, and a subject-specific intercept to account for two observations per subject (improvement from the left and right hands for each subject).

To test for a possible confounding relationship between MST performance during training and sleep-dependent memory consolidation, we estimated a linear mixed-effects model with sleep-dependent memory consolidation as the dependent variable, the mean number of correct sequences in the last three training trials and age as predictors, and a subject-specific intercept (improvement from the left and right hands for each subject).

To test for a relationship between spindle rate and sleep-dependent memory improvement in the centrotemporal region and inferior Rolandic cortex, we estimated a linear mixed-effects model with sleep-dependent memory consolidation as the dependent variable, spindle rate and age as predictors, and a subject-specific intercept. In doing so, we assumed that spindle rate in the left hemisphere was linked to memory improvement with the right hand, and spindle rate in the right hemisphere was separately linked to memory improvement with the left hand^30^.

## Data availability

Raw data were generated at Massachusetts General Hospital and the Athinoula A. Martinos Center for Biomedical Imaging. Derived data supporting the findings of this study are available from the corresponding author upon reasonable request. The code for spindle detection is available at https://github.com/Mark-Kramer/Spindle-Detector-Method

## Results

### Subject characteristics

Between January 2020 and December 2023, 23 children with RE (9 with active disease, 5F, age 6.0-12.8 years), 14 children with resolved disease (8F, age 8.8-17.8 years), and 19 control children (8F, age 6.9-18.7 years) were enrolled in this prospective study. Among children who slept, the average duration of N2 sleep per EEG recording was 22.6 min (range: 9.6-49.7 min) in active RE, 27.1 min (range: 4.1-71.3 min) in resolved RE, and 21.3 min (range: 1.8-43.7 min) in controls. We detected no significant differences in the average duration of N2 sleep between groups (one-way ANOVA, p=0.7). Subject characteristics are provided in **Table 1**.

**Table 1.** Subject characteristics.

|  | Active (n=9) | Resolved (n=14) | Control (n=19) | One-way ANOVA / Chi-square test (p) |
| --- | --- | --- | --- | --- |
| Age (years) | 10.3 (6.0-12.8) | 12.0 (8.8-17.8) | 13.1 (6.9-18.7) | 0.11 |
| Female sex (%) | 55.6 | 57.1 | 42.1 | 0.65 |
| Right handedness (%) | 100.0 | 92.9 | 94.7 | 0.73 |
| Sleep opportunity (min) | 96.1 (90.0-145.0) | 93.9 (90.0-110.0) | 92.5 (90.0-108.0) | 0.66 |

### The motor sequence task captures sleep dependent memory consolidation in children

During the nap opportunity provided, 34 children slept (7 active RE, 11 resolved RE, and 16 controls) and eight children remained awake for the duration of the opportunity (2 active RE, 3 resolved RE, and 3 controls). The duration of the nap opportunity was similar between groups (p=0.7, one-way ANOVA, **Table 1**). Among children who slept, the duration of N2 sleep was similar between groups (p=0.7, one-way ANOVA, **Table 1**).

Children who slept had a greater improvement in their performance of the MST during testing (mean 16.5%, range [-66.7, 65.6]%) than children who did not sleep (mean -5.5%, range [-44.2, 14.0]%; mean improvement between groups of 20.8%, 95% CI [7.0, 34.7]%, p=0.004; **Figure 2A-B**). This finding was qualitatively consistent in each patient group (**Figure 2C;** controls, mean improvement between groups of 23.5% (95% CI [1.5, 45.5]%, p=0.037); resolved RE, mean improvement of 34.2% (95% CI [18.3, 50.0]%, p=0.0002), active RE, mean improvement between groups of 21.5% (95% CI [-4.1, 47.2]%, p=0.094).

**Figure 2.**
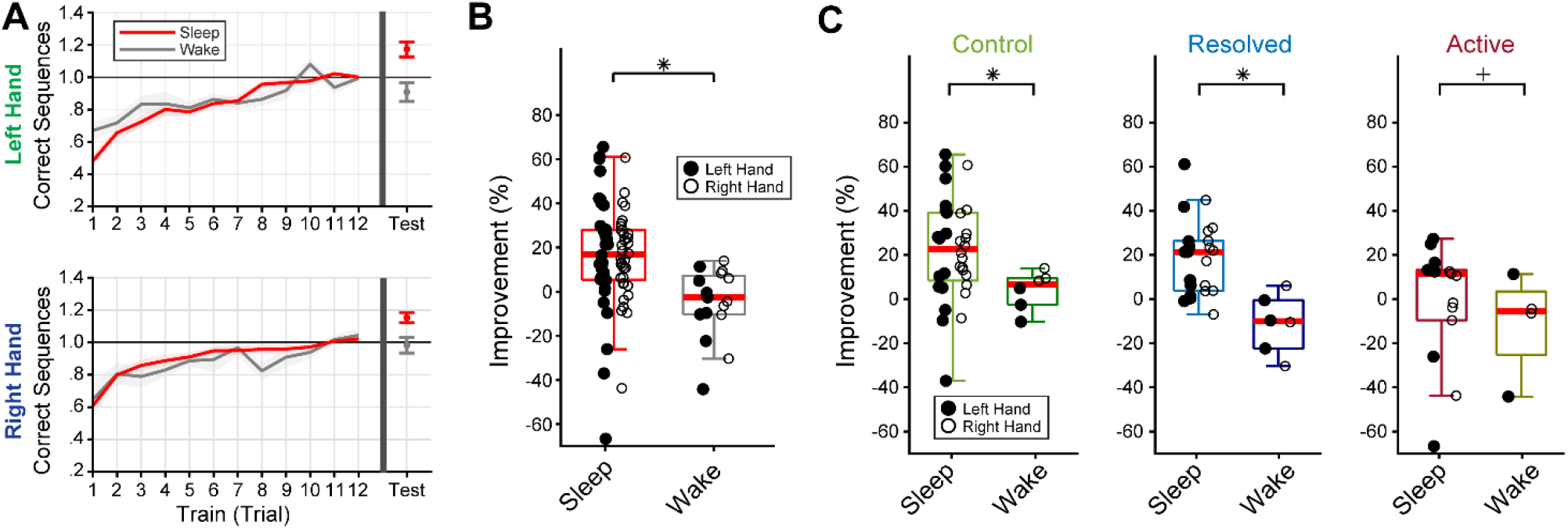
The motor sequence typing task captures sleep dependent memory consolidation. **A)** All children showed normal learning of the task during the 12 training trials (left), but children who stayed awake had decreased improvement after the nap opportunity compared to those who slept. **B)** Memory improvement of the nap opportunity is plotted for each subject, each hand. **C)** Memory improvement in children who slept compared to those who stayed awake in resolved, active RE, and control groups. *p<0.05, +p<0.1

### Children with active RE have impaired sleep-dependent memory consolidation

To evaluate for deficits in sleep-dependent memory consolidation in children with active RE, we compared the MST improvement between groups. All children demonstrated learning of the task during the training trials, with initial improvement and subsequent plateau in performance (**Figure 3A-D**). Across all groups, 4.5% of trials were excluded as outliers; there was no difference in the percentage of excluded trials between groups (controls: 4.7%; resolved RE: 4.6%; active RE: 3.9%; p=0.5, one-way ANOVA). We detected no relationship between the final performance of the task during training and the degree of sleep-dependent memory consolidation (p=0.6, linear mixed effects model). Children with active RE have decreased memory consolidation compared to control children (p=0.001, mean percentage reduction: 25.7%, 95% CI [10.3, 41.2]%) and compared to children with resolved RE (p=0.007, mean percentage reduction: 21.9%, 95% CI [6.2, 37.6]%; **Figure 3D-E**). Consistent results were observed for each hand and in all groups. Thus, children with RE have a transient deficit in sleep-dependent memory consolidation during the active phase of their disease.

**Figure 3.**
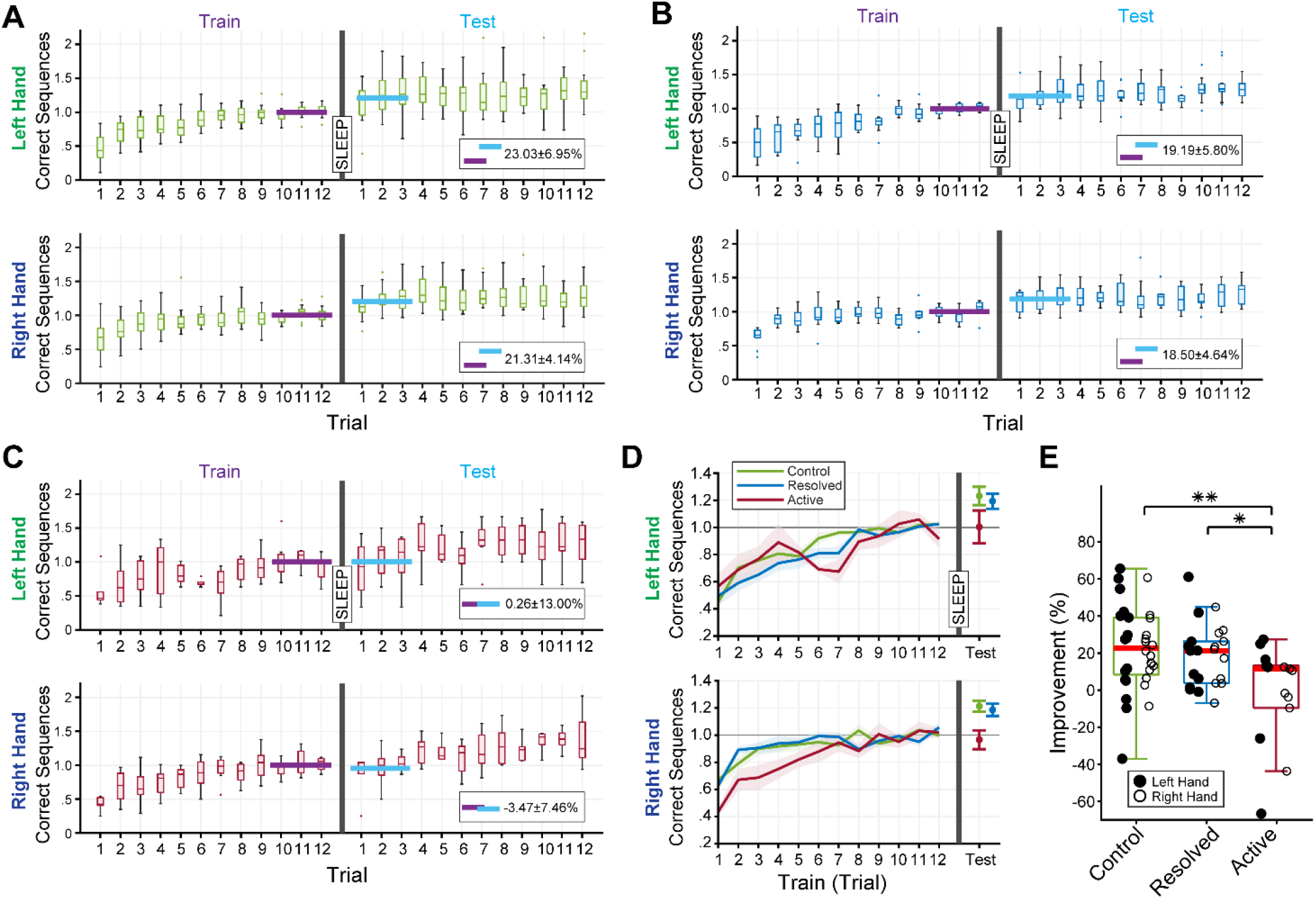
Decreased sleep-dependent memory consolidation in active RE. Box plots of correct sequences normalized by the last three trainings trials in **A)** control group, **B)** resolved RE group, and **C)** active RE group. **D-E)** Children with active RE showed normal learning of the task during the 12 training trials, but decreased sleep-dependent improvement compared to children with resolved RE and controls. **p≤0.005, *p≤0.01

### Children with active RE have decreased sleep spindles in the epileptic zone

Based on our prior findings^8^, we hypothesized that children with active RE would have decreased sleep spindles in the centrotemporal region compared to controls. Indeed, we found that children with active RE had decreased spindle rates in the central (p=0.01, mean decrease 2.4 spindles/min, 95% CI [0.6, 4.1] spindles/min) and temporal electrodes (p=0.01, mean decrease 2.8 spindles/min, 95% CI [0.7, 5.0] spindles/min) compared to the control children (**Figure 4**). Combining central and temporal channels, children with active RE had a reduced spindle rate in the centrotemporal region compared to control children (p=0.008, mean decrease 2.5 spindles/min, 95% CI [0.7, 4.4] spindles/min)^8^. We found no evidence of a difference in spindle rate between groups in any other brain region. This finding replicates our prior work^8^ in a second cohort, and we conclude that spindle rate is transiently and focally decreased in the cortical epileptic zone in children with active RE.

**Figure 4.**
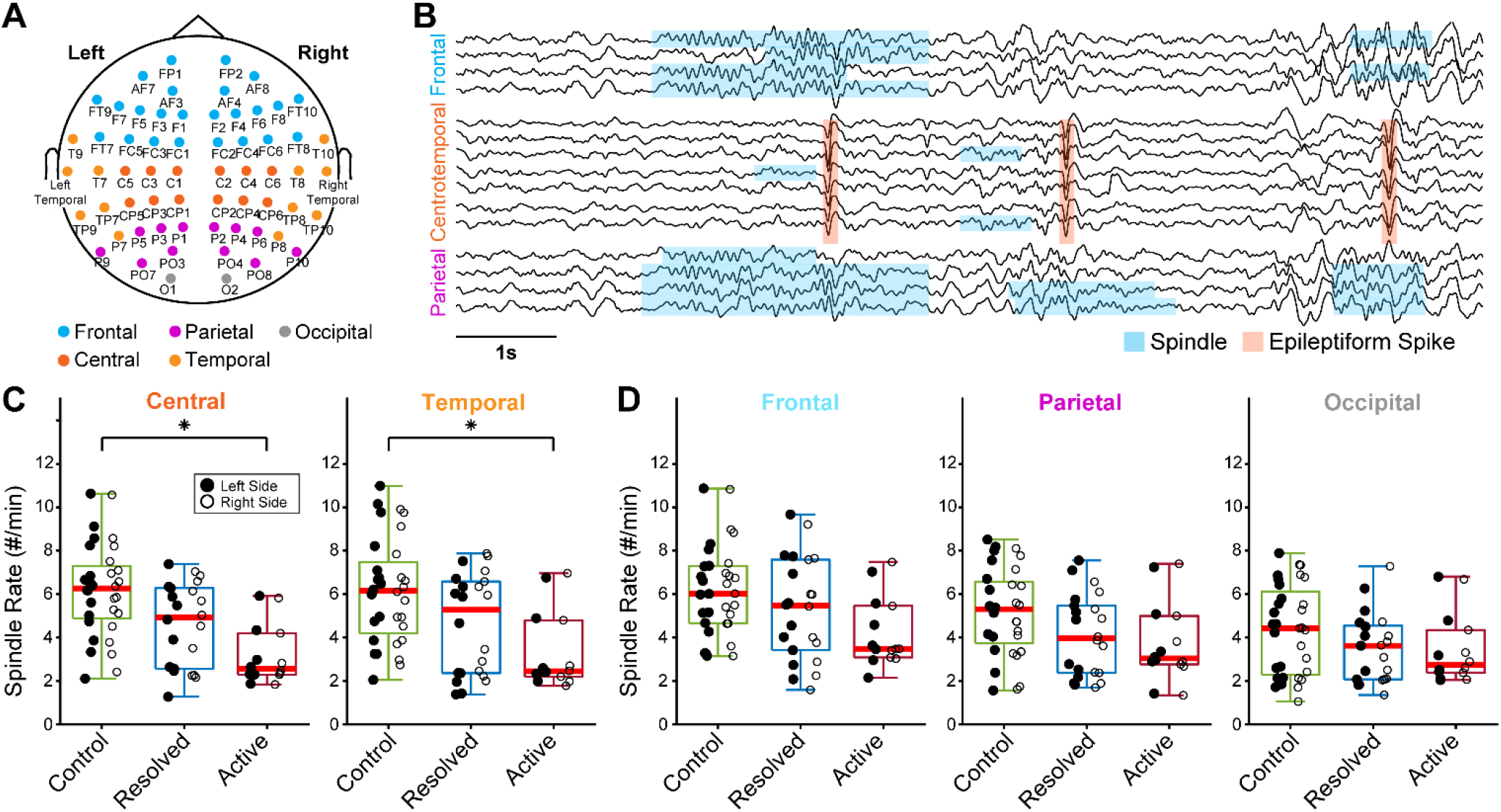
Focal sleep spindle deficit in the epileptic cortex in active RE. **A)** Illustration of electrode groups corresponding to frontal (blue), parietal (purple), occipital (gray), central (red), and temporal (orange). **B)** Example visualization showing the absence of sleep spindles (blue) in a cortical region impacted by epileptiform spikes (red). **C)** Spindle rate is decreased in the central and temporal regions in children with active RE compared to controls, * p<0.05. **D)** There is a trend toward a decrease in spindle rate in the frontal region (p=0.06) in the active RE group compared to the control group. There is no evidence of a difference in spindle rate between active RE and control groups in parietal (p=0.2) or occipital (p=0.3) regions. In each boxplot, the red line indicates the median, and the box indicate the 25th and 75th percentiles. The whiskers extend to the most extreme data points.

### Spindle rate predicts contralateral sleep-dependent memory improvement

Based on the proposed mechanistic relationship between sleep spindles and sleep-dependent memory consolidation^6^, we hypothesized that spindle rate would predict sleep-dependent memory consolidation in children as measured by performance in the contralateral hand^30^. As hypothesized, we found a positive correlation between spindle rate in the centrotemporal region and sleep-dependent memory consolidation across children (p=0.004, mean MST improvement of 3.9%, 95% CI [1.3, 6.4]%, for each unit increase in spindle rate; **Figure 5**). We found no evidence that age predicted memory consolidation (p=0.8). From these findings, we conclude that centrotemporal spindle rate predicts sleep-dependent memory consolidation in children. This finding is consistent with the proposed mechanistic relationship between sleep spindles and sleep-dependent memory consolidation.

**Figure 5.**
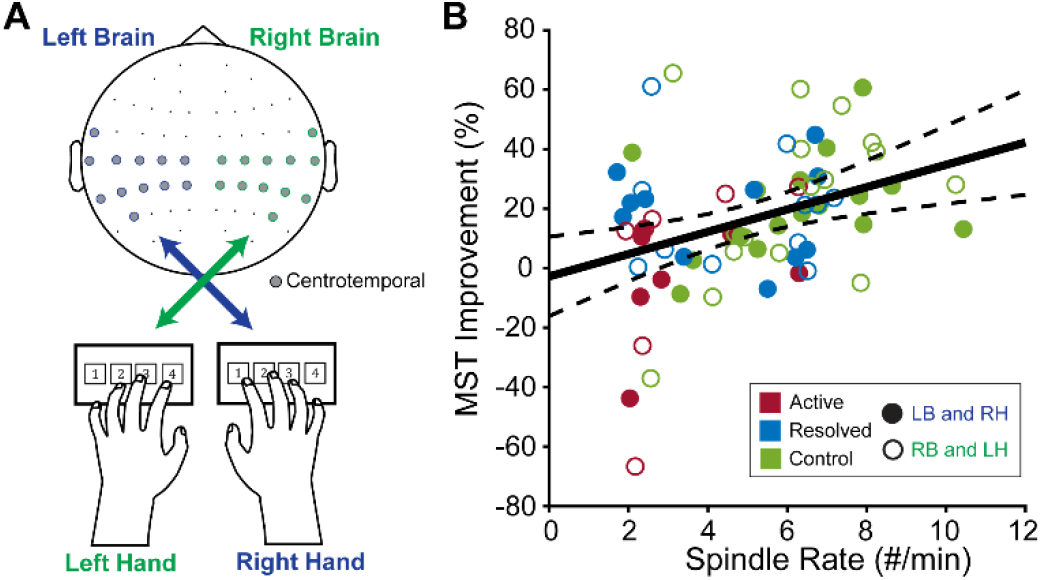
Spindle rate correlates with sleep-dependent memory improvement in the centrotemporal region. **A)** MST improvements with left and right hand are linked with the spindle rate in the contralateral hemisphere. **B)** Sleep spindle rate in the centrotemporal region positively correlates with the degree of sleep-dependent memory consolidation. Black (dashed) curves indicate estimated model fit (95% confidence interval). LB: left brain; RH: right hand; RB: right brain; LH: left hand.

### Spindle rate in the inferior Rolandic cortex is decreased in active RE and predicts sleep-dependent memory consolidation

To focus our analysis specifically on the irritative zone in RE, we performed electrical source imaging, targeting the inferior Rolandic cortex as our region of interest (see Materials and Methods). We then estimated spindle rate from this region across groups. Replicating our prior results^26^ in this new cohort, we found reduced spindle rate in the inferior Rolandic cortex in children with active RE compared to control children (p=0.027, mean decrease 1.85 spindles/min, 95% CI [0.21, 3.49], **Figure 6**). We also found that spindle rate in the inferior Rolandic cortex predicted sleep dependent memory consolidation as measured in the contralateral hand across children (p=0.014, mean MST improvement of 3.7%, 95% CI [0.8, 6.7]%, for each unit increase in spindle rate). We again found no evidence that age predicted memory consolidation (p=0.3). Consistent with results estimated from the scalp EEG, these findings support the proposed mechanistic relationship between sleep spindles and sleep-dependent memory consolidation.

**Figure 6.**
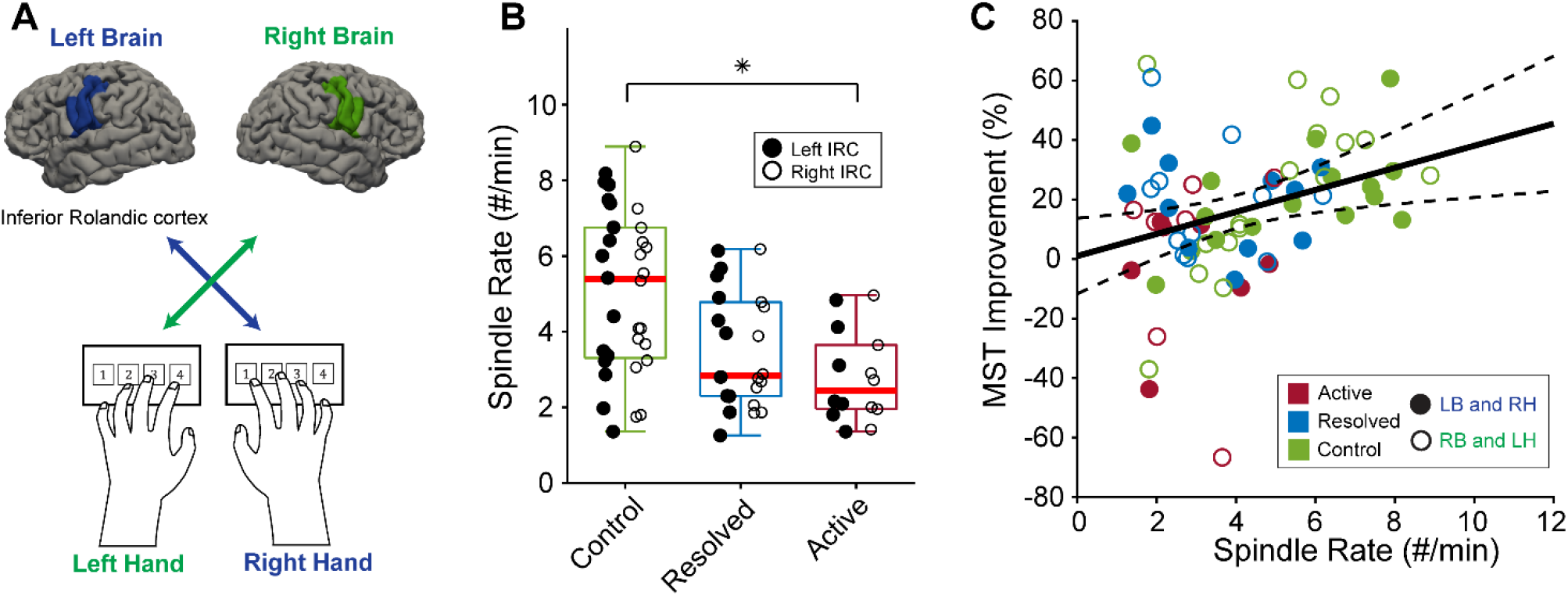
Spindle rate correlates with sleep-dependent memory improvement in inferior Rolandic cortex. **A)** MST improvements with left and right hand are linked with the spindle rate in the contralateral inferior Rolandic cortex. **B)** Children with active RE showed a reduced spindle rate, * p<0.05. **C)** Sleep spindle rate in inferior Rolandic cortex positively correlates with the degree of sleep-dependent memory consolidation. Black (dashed) curves indicate estimated model fit (95% confidence interval). IRC: Inferior Rolandic cortex ). LB: left brain; RH: right hand; RB: right brain; LH: left hand.

## 4. Discussion

Sleep spindles are prominent thalamocortical brain oscillations in the sleep EEG that have been mechanistically linked to essential sleep-dependent cognitive processes in animal models^7,31,32^ and healthy controls^13,19,22,33^. Here we identify a deficit in sleep-dependent memory consolidation in Rolandic Epilepsy, the most common focal developmental epilepsy in childhood. We find that a transient, focal deficit in sleep spindle rate correlates with a transient deficit in sleep-dependent memory consolidation. Further, that sleep spindle rate positive predicts sleep dependent memory consolidation in RE and control children.

Increasing evidence suggests that sleep spindles are a key oscillatory mechanism required for off-line memory consolidation during N2 sleep^34^. Sleep spindles are generated by GABAergic neurons in the thalamic reticular nucleus and coordinated by well-delineated thalamocortical circuits. Spindle rate is higher in cortical regions that are linked to prior learning experiences^15^, and spindles accompany the cortical reactivation patterns of memory replay during sleep^20^. In rodent models, calcium recordings during sleep suggest that sleep spindles gate dendritic calcium shifts required for synaptic plasticity. Spindle oscillations also coordinate hippocampal sharp wave ripple oscillations that reflect neuronal replay in rodents^7,31,35^ and in humans^36^. In healthy adults, the relationship between sleep spindles and sleep-dependent memory consolidation is well-established for both procedural^13,14,22,30,33^ and declarative^15,16,21^ tasks. In healthy children, sleep spindle rate, not sleep duration^37^, predicts general cognitive abilities^18,38^ and sleep-dependent memory consolidation for declarative^18,19^ and procedural^39^ tasks. Sleep spindle rate has been reported to be decreased in autism, developmental delay, and attention deficit hyperactivity disorder^40,41^ and correlate with cognitive abilities in children with dyslexia^42^ and epilepsy^8^. Critically, several studies have found that medications and interventions that increase sleep spindles improve memory consolidation, and medications that disrupt spindles impair memory consolidation^9,43-45^, suggesting a mechanistic link between sleep spindles and memory performance. Further supporting sleep spindles as generalizable biomarker for sleep-dependent memory processes across ages and conditions, we find that focal spindle rate predicts sleep dependent memory consolidation on a motor procedural task in healthy children and children with focal epilepsy. The motor sequence typing task involves learning and typing a 5-digit sequence as quickly and accurately as possible during a training period, and again during a testing period, separated by wakefulness or sleep. This task has been validated to capture sleep-dependent memory consolidation in healthy adults^22^, adults with schizophrenia and adults with epilepsy^46^. In healthy children, one study found robust sleep-dependent improvements in the motor sequence task in 9-11 years old children while two studies in 6-11 years old children failed to detect sleep-dependent improvements^37,47^. In this study, we analyzed children 6-18 years old from three groups – children with active RE, in remission, and control subjects – and found greater improvements in the performance of the motor sequence task in children who slept compared to those who did not, and a positive relationship between the degree of consolidation and sleep spindle rate. In our cohort, we did not find a relationship between the degree of consolidation and age. Our use of a short nap opportunity, where the state of consciousness was observed continuously by EEG, may have afforded a more sensitive detection of sleep than self-report used in prior studies. These findings support the motor sequence task as a sensitive measure of sleep-dependent memory consolidation in school-age children.

Although our sample size was relatively small, our findings of a focal spindle deficit in RE replicate results from a prior study^8^, increasing the rigor of this finding. Here, we also used short naps instead of a full night of sleep, as prior work has demonstrated that nocturnal spindle rate can be reliably estimated from daytime naps in adults^48^ and predict several measures of cognitive ability in children^8,49^. However, a full night could capture further sleep dynamics, such as alterations in sleep homeostasis^50^, which were not evaluated here.

In conclusion, children with active RE have deficits in sleep-dependent memory consolidation that can be predicted by non-invasive measures of sleep spindle rate. This work supports sleep spindles as a mechanistic biomarker for memory consolidation in children with RE and controls, and implicates disruption of this rhythm in the cognitive symptoms observed in RE.

## Author Contribution

H.K and C.J.C planned the study, analyzed the data; H.K. wrote the first draft of the manuscript; H.K, D.M.C, E.A.K, S.K.G, J.F.H, E.D.B, K.G.W, W.S and C.J.C collected data, wrote the manuscript; M.A.K, D.S.M contributed to developing analysis methods and provided feedback; C.J.C supervised the study.

## Acknowledgments

This work was supported by NINDS R01NS115868.

